# ALK signaling drives tumorigenicity and chemoresistance of pancreatic ductal adenocarcinoma cells

**DOI:** 10.1101/2022.08.29.505637

**Authors:** Beatriz Parejo-Alonso, Alba Royo-García, Pilar Espiau-Romera, Sarah Courtois, Álvaro Curiel-García, Sladjana Zagorac, Isabel Villaoslada, Kenneth P. Olive, Christopher Heeschen, Patricia Sancho

**Affiliations:** Instituto de Investigación Sanitaria Aragón (IIS Aragón), Zaragoza, Spain; Department of Medicine, Division of Digestive Liver Diseases and Herbert Irving Comprehensive Cancer Center, Columbia University Irving Medical Center, New York, New York, USA; Centre for Stem Cells in Cancer & Ageing (Barts Cancer Institute), London, UK; Center for Single-Cell Omics and Key Laboratory of Oncogenes and Related Genes, Shanghai Jiao Tong University School of Medicine, China; Pancreatic Cancer Heterogeneity, Candiolo Cancer Institute - FPO – IRCCS, Candiolo (Torino), Italy

**Keywords:** Pancreatic Ductal Adenocarcinoma, ALK, Receptor Tyrosine Kinases, Cancer Stem Cells, Chemoresistance

## Abstract

Pancreatic ductal adenocarcinoma (PDAC) is an extremely aggressive disease characterized by its metastatic potential and chemoresistance. These traits are partially attributable to the highly tumorigenic pancreatic cancer stem cells (PaCSCs). Interestingly, these cells show unique features in order to sustain their identity and functionality, some of them amenable for therapeutic intervention. Screening of phospho-receptor tyrosine kinases revealed that PaCSCs harbored increased activation of anaplastic lymphoma kinase (ALK). We subsequently demonstrated that oncogenic ALK signaling drives tumorigenicity in PDAC patient-derived xenografts (PDXs) by promoting stemness through ligand-dependent activation. Indeed, the ALK ligands midkine (MDK) or pleiotrophin (PTN) increased self-renewal, clonogenicity and CSC frequency in several *in vitro* local and metastatic PDX models. Conversely, treatment with the clinically-approved ALK inhibitors Crizotinib and Ensartinib decreased CSC content and functionality *in vitro* and *in vivo*, by inducing cell death. Strikingly, ALK inhibitors sensitized chemoresistant PaCSCs to Gemcitabine, as the most used chemotherapeutic agent for PDAC treatment. Consequently, ALK inhibition delayed tumor relapse after chemotherapy *in vivo* by effectively decreasing the content of PaCSCs. In summary, our results demonstrate that targeting the MDK/PTN-ALK axis with clinically-approved inhibitors impairs *in vivo* tumorigenicity and chemoresistance in PDAC suggesting a new treatment approach to improve the long-term survival of PDAC patients.

**GRAPHICAL ABSTRACT:** 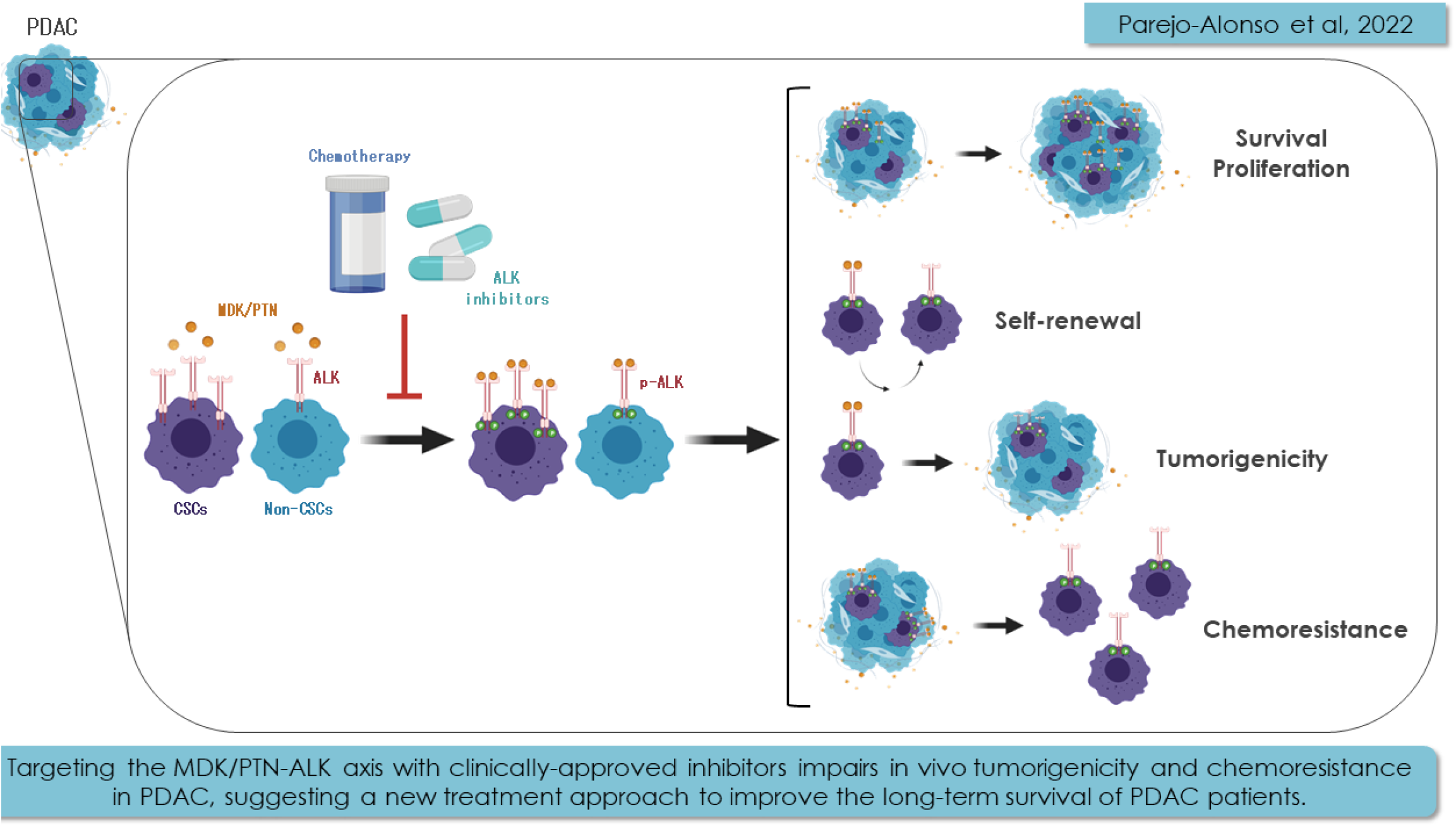

## INTRODUCTION

Pancreatic ductal adenocarcinoma (PDAC) is the most represented pancreatic cancer, with an increasing incidence and an extremely poor five-year overall survival of less than 9%^[1,2]^. Despite PDAC low incidence rate compared to other malignancies, its mortality rate continues to rise, potentially making it one of the three most deadly cancers by 2030^[3,4]^. Importantly, PDAC aggressiveness mainly relies on two key factors: delayed detection, since symptoms are mild and/or unspecific even at advanced disease stage, and its intrinsic resistance to current treatment regimens^[1]^. Regarding the latter, PDAC is a considerably heterogeneous disease organized hierarchically, with mounting evidence of a small but unique subpopulation of cancer cells with self-renewal capacity and tumor-initiating properties. These pancreatic cancer stem cells (PaCSCs) are capable of symmetrical and asymmetrical divisions, the former giving rise to identical CSCs to perpetuate its lineage, and the latter to differentiated progenies that form the bulk of the tumor^[5,6]^. As such, PaCSCs have the capacity to initiate and sustain tumor growth, in addition to promoting recurrence after treatment due to their intrinsic chemoresistance^[7]^. Therefore, new treatment strategies for pancreatic cancer are urgently needed. One of the most explored avenues to design new treatment strategies is the inhibition of tyrosine kinases (TKs), which are hyperactivated in many cancer types and control basic functions such as cell differentiation, proliferation and metabolism^[8,9]^. Notably, several TK inhibitors (TKi) are currently under evaluation in clinical trials for PDAC, including Erlotinib (EGFR inhibitor, phase III) or Masitinib (c-kit/PDGFR inhibitor, phase III)^[10]^. Interestingly, several receptor TKs (RTKs) have been shown to promote stemness in different tumor types, such as EGFR in breast cancer^[11,12]^, FGFR in prostate cancer^[13]^, EphR in brain tumors^[14,15]^, melanoma^[16]^ and lung cancer^[17]^, as well as the non-receptor TK SRC in pancreatic cancer^[18]^. Hence, RTKs may also be potential candidates for targeting PaCSCs and thereby improve patient long-term survival.

Here, we show now that the RTK anaplastic lymphoma kinase (ALK) and its downstream signaling pathways are overactivated in PaCSCs. Mechanistically, we demonstrate that ALK can be exogenously activated by midkine (MDK) and drives essential stemness functions such as self-renewal and tumorigenicity. Crucially, we found that pathway blockade with clinically-approved ALK inhibitors rescue Gemcitabine resistance in PDAC, thereby providing a new perspective for a more effective treatment regimen against this deadly disease.

## Experimental Procedures

### Cell Culture

#### Patient-Derived Xenografts (PDXs) and Circulating Tumor Cells (CTCs)

PDX185, PDX215, PDX253 and PDX354 were obtained through the Biobank of the Spanish National Cancer Research Center (CNIO, Madrid, Spain; CNIO20-027). Tumor pieces underwent several amplification passages in mice prior to establishing primary cultures. Tumor dissociation and establishment of *in vitro* primary cultures was performed as previously described^[19]^. The metastatic model CTCA was established from circulating tumor cells and obtained through the Barts Pancreas Tissue Bank of the Barts Cancer Institute (https://www.bartspancreastissuebank.org.uk/; BCI, London, United Kingdom; 2019/02/IISA/PS/E/Cell-cultures). Cells were submitted to a maximum of 15 passages in complete RPMI 1640 GlutaMAX™ medium supplemented with 10% FBS and 50 U/mL penicillin/streptomycin (all from Gibco, Life Technologies, Carlsbad, CA, USA). For experiments, the medium was changed to DMEM/F-12 GlutaMAX™ supplemented with 2% B27 (Gibco), 50 U/mL penicillin/streptomycin, and 20 ng/mL FGF-basic (Pan-Biotech, Aidenbach, Germany). All PDXs were grown in a humidified incubator at 37ºC with 5% CO_2_ and regularly tested for mycoplasma at the Technical and Scientific Services Unit from the Health Sciences Institute of Aragón (IACS).

#### Cell Lines

AsPC1, BxPC3, MiaPaCa2, Panc1 and Su8686 were purchased from the American Type Culture Collection (ATCC, Manassas, VA, USA). Cells were submitted to a maximum of 30 passages in the same conditions as the primary cultures described above.

#### Spheroids

Cells were cultured in anchorage-independent conditions at 10^5^ cells/mL with complete DMEM/F-12 GlutaMAX™ medium. First generation spheroids were grown up to seven days, dissociated with trypsin (Corning, Oneonta, NY, USA) and regrown at 10^5^ cells/mL for five more days. Flasks were coated with 10% poly(2-hydroxyethyl methacrylate) (Sigma-Aldrich, Saint Louis, MO, USA) in 96% ethanol and left at 37ºC until all the liquid was evaporated. Flasks were rinsed once with 1X PBS prior to their utilization.

### *In vitro* Treatments

*ALK inhibitors*: Crizotinib and Ensartinib were purchased from Selleckchem (Munich, Germany) and dissolved in DMSO (Sigma-Aldrich) following the manufacturer’s instructions. Cells were treated for 24 to 72 hours at concentrations ranging 0.5 to 10 µM, with DMSO compensation when needed.

#### Chemotherapy

Gemcitabine 0.9% sodium chloride (Eli Lilly and Company, IN, USA) was used at concentrations ranging from 1 to 750 nM for 24 to 72 hours.

### Flow cytometry

#### Apoptosis

Cell pellets were rinsed once with 1X PBS and incubated on ice for 15 minutes in 2% FBS-0.5% BSA-1X PBS blocking solution. PE-conjugated CD133 antibody or the corresponding control immunoglobulin G1 were added at 1:400 in blocking solution. Cells were stained on ice for 30 minutes and protected from light. Then, the antibody or IgG1 excess was rinsed and pellets were resuspended in APC-conjugated Annexin V at 1:20 in Annexin V buffer solution plus Zombie Violet dye at 1:400 (all antibodies and probes are from Biolegend, San Diego, CA, USA). Samples were transferred into FACS tubes and incubated for 20 minutes at room temperature protected from light prior to their analysis by FACS Canto II (BD, Franklin Lakes, NJ, USA). Flowing 2 software (Turku Bioscience Centre, Turku, Finland) was used for data analysis.

#### Fluorescence Activated Cell Sorting (FACS)

Cells were blocked and stained for CD133 as described above. For autofluorescence sorting, a previously described protocol was followed^[20]^. After staining, pellets were resuspended in Zombie Violet dye at 1:400 in 1X PBS and incubated for 20 minutes at room temperature protected from light. Viable cells corresponding to CD133 or autofluorescence negative and positive populations were sorted using the SH800S Cell Sorter (Sony Biotechnology, San José, CA, USA) and collected into 5 mL tubes containing complete RPMI medium. Pellets were stored at −80ºC for further processing.

### Proteome Profiler™ Array

Cells were sorted by autofluorescence and CD133 expression using the SH800S Cell Sorter as described above and pellets were lysed according to manufacturer’s instructions. The samples were further processed following the Human Phospho-Receptor Tyrosine Kinase Kit (R&D Systems Europe, Ltd., Abingdon OX14 3NB, UK) manufacturer’s instructions. Pierce™ ECL Western Blotting Substrate was used to detect protein-antibody complexes prior to visualization on CL-X Posure™ films (both from ThermoFisher Scientific, Waltham, MA, USA). The resulting dots were analyzed using ImageJ software (National Institutes of Health, Bethesda, MD, USA).

### Western Blot

Cell pellets were lysed in RIPA buffer (Sigma-Aldrich) plus protease and phosphatase inhibitors (both from Alfa Aesar, ThermoFisher Scientific). After extraction, proteins were quantified using the Pierce™ BCA Protein Assay Kit (ThermoFisher Scientific). Proteins were separated in Novex™ WedgeWell™ 10% Tris-Glycine precast gels using BenchMark™ pre-stained protein ladder (both from Invitrogen, Carlsbad, CA, USA) and transferred into PVDF membranes (ThermoFisher Scientific). Membranes were blocked in 5% BSA-1X PBS-0.1% Tween 20 (ThermoFisher Scientific) for 1 hour at room temperature and incubated overnight at 4ºC with the following primary antibodies: ALK (1:1000), pALK (Tyr1604, 1:1000), ERK 1/2 (1:3000) (all from Cell Signaling Technology, Danvers, MA, USA), pERK 1/2 (T202-Y204, 1:3000, Abgent, San Diego, CA, USA), and β-actin as loading control (clone AC-74, 1:10 000, Sigma-Aldrich). After several washes with 1X PBS-0.5% Tween 20, membranes were incubated with peroxidase-conjugated goat anti-rabbit (1:5000) or goat anti-mouse (1:50000) (both from Invitrogen). Pierce™ ECL Western Blotting Substrate was used to detect protein-antibody complexes prior to visualization on CL-X Posure™ films. Band intensities were analyzed using ImageJ software. Likewise, protein from sorted CD133 and autofluorescence cells extracted for the RTK array were separated, transferred and visualized as described for normal Western Blot.

### Real Time Quantitative Polymerase Chain Reaction (RTqPCR)

Cell pellets were homogenized in TRIzol reagent (Invitrogen) and RNA was extracted according to manufacturer’s instructions and quantified using Nanodrop™ 2000 (ThermoFisher Scientific). 1 µg of RNA was retrotranscribed into cDNA using Maxima H minus cDNA synthesis Master Mix with dsDNase kit (ThermoFisher Scientific). RTqPCR was performed using PowerUp™ SYBR Green Master Mix (Applied Biosystems, ThermoFisher Scientific) according to manufacturer’s instructions. The primers used are listed below. *HPRT* was used as endogenous housekeeping gene.

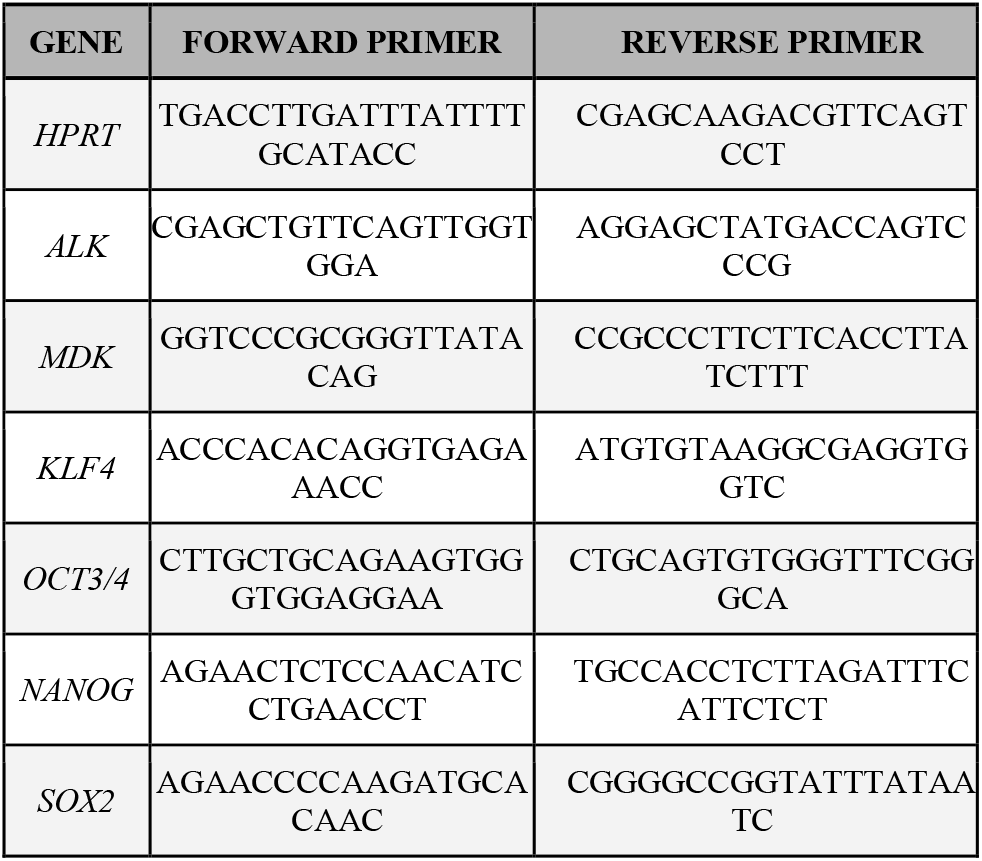

### Bioinformatic Analyses

Expression data from human PDAC tissue and normal pancreatic tissue were analyzed using the webserver GEPIA2 (TCGA and the GTEx project databases; http://gepia2.cancer-pku.cn/)^[21]^. The Pearson correlation coefficient was calculated to study the association of the individual genes corresponding to ALK ligands with a stemness signature defined by the combined expression of the pluripotency-related genes *KLF4, OCT3/4, NANOG* and *SOX2*. A publicly available human single-cell RNA sequencing (scRNAseq) dataset^[35]^ was used to investigate the expression of *ALK, MDK* and *PTN* genes. Raw counts were obtained from the Genome Sequence Archive (#CRA001160). Raw counts from tumor samples were filtered excluding low-quality cells and normalized using CPM (counts per million). Transform function was used to obtain the final gene expression matrix. Cell types identified in the reference papers were also used to calculate and plot the expression and percentage of cells in each group for *ALK, MDK* and *PTN* genes using ggplot2 package. All analyses were performed using R v.4.2.1. The PDAC samples of the TCGA dataset were classified into high and low *ALK* expression and compared in gene set enrichment analyses (GSEA). The GSEA module of the GenePattern suite from the Broad Institute was used with 1000 permutations and FDR < 25% was considered statistically significant. The signatures for stemness and *ALK* overexpression (OE) were previously described in Ai *et al*.^[22]^ and Mazzeschi *et al*.^[23]^, respectively. *ALK* mutational status was assessed using the webserver cBioPortal (Pancreas UTSW, Pancreas TCGA PanCan 2018, Pancreas TCGA, Pancreas ICGC and Pancreas QCMG 2016 project datasets; https://www.cbioportal.org.)[24].

### Enzyme-Linked Immunosorbent Assay (ELISA)

MDK levels in supernatants from cell cultures and fresh tumor pieces, as well as plasma from mice bearing orthotopic tumors were determined using the MDK DuoSet ELISA kit (R&D systems, Minneapolis, MN, USA) as per manufacturer’s instructions. For MDK determination in tumor pieces, freshly extracted subcutaneous or orthotopic tumors were minced, and pieces of around 1 mm^3^ were incubated for 24 hours in 1 mL of complete DMEM/F12 medium.

### Sphere Formation Assay (SFA)

10^4^ cells were seeded in triplicate in complete DMEM/F-12 medium using polyhema-coated 24-well plates in the presence of different treatments. When indicated, cells were pre-treated in adherence for 48 hours in complete DMEM/F-12 medium prior to being seeded without treatments in anchorage-independent conditions as described above. In both cases, the spheroids were counted after seven days using an inverted microscope at 20X magnification.

### Colony Formation Assay (CFA)

Cells were seeded in complete RPMI medium in 6-well plates at a density of 500 or 1000 cells/well. After 24 hours, treatments were added in complete DMEM/F-12 medium. Medium and treatments were changed every seven days. After 21 days, colonies were stained with crystal violet dye (Acros Organics, ThermoFisher Scientific). Colonies were then counted manually, dissolved in 1% sodium dodecyl sulfate (SDS, ThermoFisher Scientific) and the absorbance at 590 nm was read using the plate reader Synergy HT (BioTek Instruments, Santa Clara, CA, USA).

### Extreme Limiting Dilution Assay (ELDA)

#### In vitro ELDA

10^3^ cells per condition were mixed with DMEM/F-12 medium plus treatments and serial dilutions were then seeded in sextuplicate in polyhema-coated 96-well plates. After seven days, the presence or absence of, at least, one spheroid was assessed using an inverted microscope. Further analysis was done by the Walter+Eliza Hall Bioinformatics online tool for ELDA analysis (http://bioinf.wehi.edu.au/software/elda/)^[25]^.

#### In vivo ELDA (Tumorigenicity Assay)

Cells were pre-treated *in vitro* for 48 hours. Two cell densities (10^4^ and 10^3^) diluted in 50:50 complete DMEM/F-12 medium:Matrigel™ (Corning) were subcutaneously injected into both the top and bottom flanks of six weeks-old Foxn1^nu^ nude mice of both sexes (n=4 mice per group, n=8 injections per group). Tumor size was monitored once a week using a caliper and volumes were calculated using the formula (length*width^2^)/2. After six weeks, when control mice had reached humane endpoint criteria, mice were euthanized, tumors were collected and pictures were taken. The number of tumors at end point was analyzed using the Walter+Eliza Hall Bioinformatics online tool, considering tumors >50 mm^3^ that were growing for 3 weeks in a row, the rest were excluded from the analysis. Tumors corresponding to the injections with 10^4^ cells from PDX354 were dissociated and stained with EpCAM-FITC, CD133-PE and CD44-APC antibodies for FACS analysis as described above. Mice were housed according to institutional guidelines and all experimental procedures were performed in compliance with the institutional guidelines for the welfare of experimental animals as approved by the Universidad of Zaragoza Ethics Committee (CEICA PI22/17) and in accordance with the guidelines for Ethical Conduct in the Care and Use of Animals as stated in The International Guiding Principles for Biomedical Research involving Animals, developed by the Council for International Organizations of Medical Sciences (CIOMS).

### Viability assay

Cells were seeded in triplicate in 96-well plates 24 hours before treatment. At zero and 72 hours, medium was discarded and Resazurin (Alfa Aesar) was added to the cells at 10 µM in 1X PBS and incubated for one hour in a humidified incubator at 37ºC with 5% CO_2_. Fluorescence was then read according to manufacturer’s instructions by using a Synergy HT plate reader. The IC_50_ was calculated using GraphPad Prism 8.

### MultiTox-Fluor Multiplex Cytotoxicity Assay

Cells were seeded in triplicate in 96-well plates 24 hours before treatment. At zero and 72 hours, assay was performed by incubating with Multitox reagents (Promega, Madison, WI, USA) and fluorescence was then read according to manufacturer’s instructions by using a Synergy HT plate reader.

### Proliferation assay

After Resazurin or MultiTox technique, the cells were rinsed once with 1X PBS and incubated for 30 minutes with crystal violet. The plates were rinsed carefully with tap water and dried for at least 24 hours. After dissolution in 1% SDS, the absorbance at 590 nm was read using a Synergy HT plate reader. The IC_50_ was calculated using GraphPad Prism 8.

#### *In Vivo* Treatment

Tumor pieces of about 15 mm^3^ were soaked in Matrigel™ prior subcutaneous implantation in both flanks of six weeks-old Foxn1^nu^ nude female mice (n=4 mice per group, n=8 implants per group). When tumor size was about 300 mm^3^, mice were treated with one cycle of chemotherapy as follows: 30 mg/kg Abraxane (i.v.) twice a week plus 70 mg/kg Gemcitabine (i.p) once a week during three weeks and one week of rest. After the chemotherapy cycle, mice were randomized and treated with 25 mg/kg Crizotinib or the corresponding dose of vehicle (hydroxypropyl methyl cellulose, Sigma Aldrich) (oral gavage) twice a day until endpoint. Tumor size was monitored twice a week using a caliper and volumes were calculated using the formula (length*width^2^)/2. After 10.5 weeks, when control tumors had reached humane endpoint criteria, mice were euthanized, tumors were collected and weighted and pictures were taken. A small piece of the tumors was processed for RNA to assess pluripotency gene expression by RTqPCR as described above. The rest of the tumors was dissociated as previously reported^[19]^ and stained with EpCAM-FITC, CD133-PE and CD44-APC antibodies for FACS analysis as described above. Mice weight was monitored twice a week throughout the experiment in order to detect any sign of toxicity. Mice were housed according to institutional guidelines and all experimental procedures were performed in compliance with the institutional guidelines for the welfare of experimental animals as approved by the Universidad of Zaragoza Ethics Committee (CEICA PI22/17), and in accordance with the guidelines for Ethical Conduct in the Care and Use of Animals as stated in The International Guiding Principles for Biomedical Research involving Animals, developed by the Council for International Organizations of Medical Sciences (CIOMS).

### Statistical Analysis

Data are represented as mean ± SEM of, at least, three independent experiments unless otherwise specified. Data were analyzed using GraphPad Prism 8. Student’s *t*-test or Mann-Whitney test were performed for two-group comparisons, while one-way ANalysis Of VAriance (ANOVA) or Kruskal-Wallis tests were performed for multiple group comparisons. Differences were considered significant when *p* < 0.05.

### RESULTS

#### ALK receptor expression and activation are linked to stemness in PDAC patients

In order to identify pharmacologically targetable RTKs specifically activated in PaCSCs, we ran a series of RTK arrays using different CSC-enriching conditions *versus* differentiated cells (autofluorescence, **Fig. 1A, top and middle panel;** CD133, **Fig. 1A, bottom panel**). Among the differentially phosphorylated RTKs, the receptor anaplastic lymphoma kinase (ALK) showed the most consistent upregulation in both CSC-enriching conditions across the panel of tested PDXs (**Fig. 1A and S1A**). ALK receptor was shown to be aberrantly expressed and/or activated in several cancer types^[26–29]^, but its expression had been previously reported to be very low or even absent in PDAC tissues^[30]^. For this reason, we decided to verify its expression by western blot in a panel of PDAC patient-derived xenografts (PDXs) and established cell lines. Surprisingly, all of them showed considerable ALK expression and phosphorylation (**Fig. S1B**). We further validated our initial findings by western blot (**Fig. 1B**), where CSC-enriching conditions such as Fluo^+^, CD133^+^ and spheroids showed further enhanced ALK phosphorylation. Notably, CSC-enriching samples also showed increased ALK mRNA (**Fig. S1C**) and protein expression (**Fig. 1B, 1C**).

**Figure 1.**
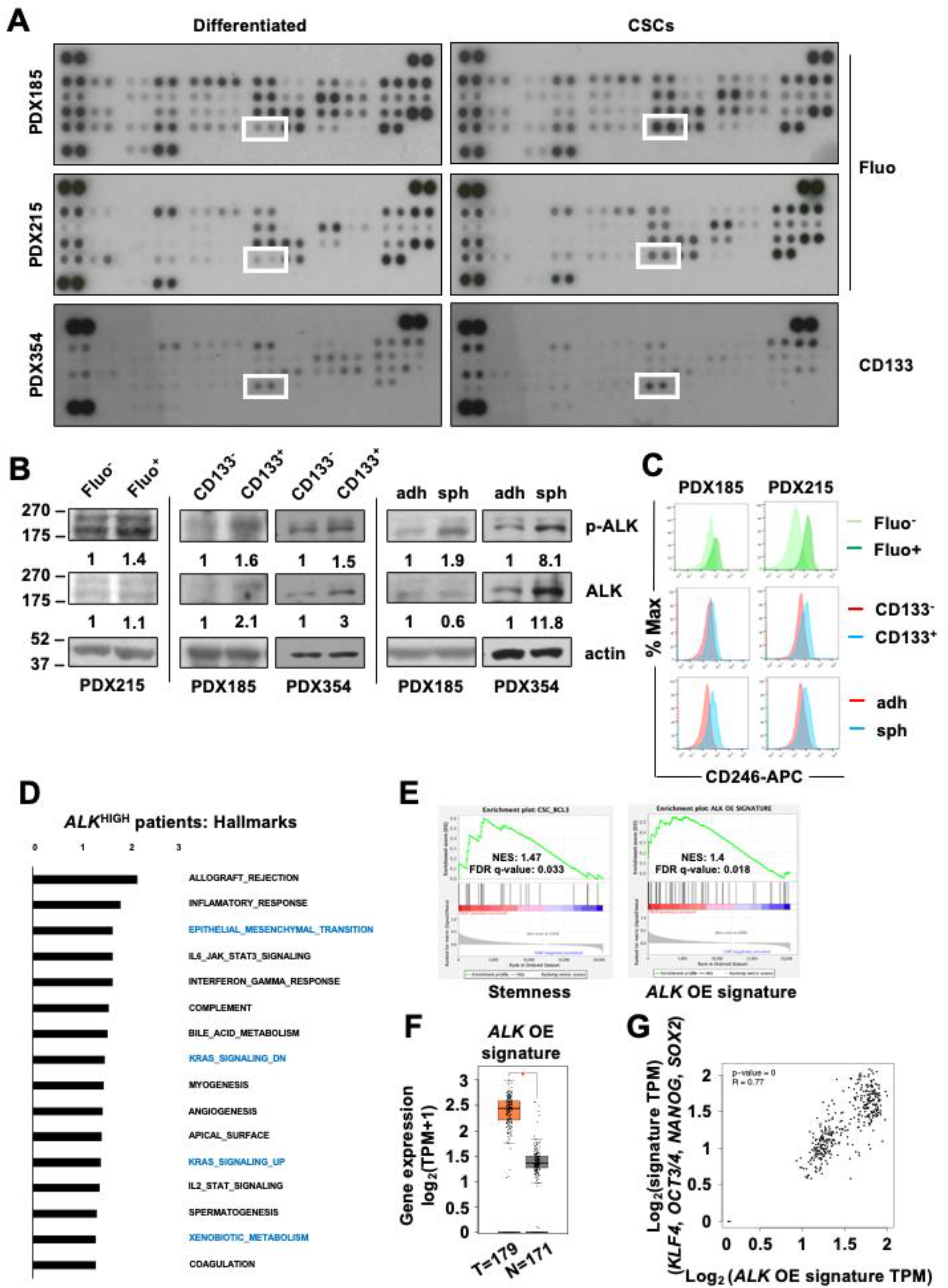
ALK is preferentially activated and overexpressed in PaCSCs. **A)** Proteome Profiler Human Phospho-RTK Array in cells sorted by autofluorescence (Fluo) and CD133 content as CSC-enriching conditions, for the indicated PDXs. Dots corresponding to pALK are indicated with a white square. **B)** Western blot of cell lysates from different CSC settings (sorted cells by autofluorescence (Fluo) and CD133 content; cells grown in adherent (adh) *versus* anchorage-independent conditions as spheroids (sph)). The numbers represent the quantification of the band intensity of each protein normalized by actin, shown as the fold change of each differentiated cell condition. **C)** Flow cytometry histograms of CD246 (ALK) expression (percentage of maximal fluorescence) in the CSC-enriching conditions shown in B. **D)** Gene set enrichment analysis comparing the top 50% *ALK* expression group (*ALK*^HIGH^) with the bottom 50% expression group in the TCGA data series. NES (normalized enrichment score) values of the Hallmark gene sets meeting the significance criteria: nominal *p*-value of <0.05, FDR <25%. **E)** Enrichment plot of stemness (left panel) and *ALK* overexpression (OE) (right panel) signatures in *ALK*^HIGH^ *versus ALK*^LOW^. **F)** Transcriptomic bioinformatic analyses of *ALK* OE signature comparing normal (N) to PDAC (T) human tissues from TCGA and GTEx datasets. **G)** Correlation of *ALK* OE signature with a stemness gene signature composed by *KLF4, OCT3/4, NANOG* and *SOX2* comparing normal (N) to PDAC (T) human tissues from TCGA and GTEx datasets. TPM: transcripts per million.s

As mentioned above, *ALK* expression is low in PDAC tissues and chromosomal translocations are very rare^[30–32]^. Indeed, although we detected a positive tendency, bioinformatic analyses showed no significant differences in *ALK* expression in PDAC patients from The Cancer Genome Atlas (TCGA) and Genotype-Tissue Expression (GTEx) datasets, as compared to healthy pancreatic tissue (**Fig. S1D**) and revealed a low percentage of genetic alterations in this gene (**Fig. S1E**). Despite its low expression, we were able to classify these PDAC patients into high and low *ALK* expression groups for gene set enrichment analysis (GSEA). Interestingly, while *ALK*^LOW^ patients did not show any enrichment, patients with higher *ALK* expression exhibited significant enrichment of pathways related to CSC properties and functionality such as epithelial-to-mesenchymal transition (EMT) and xenobiotic metabolism (**Fig. 1D**), as well as a stemness signature previously described in PDAC (**Fig. 1E**)^[22]^. On the other hand, besides pathways related to ALK downstream signaling such as K-Ras or JAK/STAT, *ALK*^HIGH^ samples also showed enrichment of an *ALK* overexpression (OE) gene signature described in breast cancer cells^[23]^ (**Fig. 1E**). Interestingly, this *ALK* OE signature was significantly overexpressed in human PDAC samples (**Fig. 1F**) and correlated with the expression of our validated set of pluripotency genes (*KLF4, NANOG, OCT3/4* and *SOX2*)^[33]^ (**Fig. 1G**).

In summary, our results indicate that ALK expression and activation is enhanced in PaCSC from different PDXs, and its expression is linked to stemness and CSC-related pathways in human PDAC samples.

#### Ligand-dependent activation of ALK contributes to PDAC stemness

Several molecules have been proposed as ALK activators, including midkine (MDK), pleiotrophin (PTN), and family with sequence similarity 150 members A (FAM150A) and B (FAM150B)^[27,34]^. Interestingly, bioinformatic analyses revealed that *MDK* and *PTN*, but not *FAM150A* nor *FAM150B*, were significantly overexpressed in human PDAC samples when compared to normal pancreas (**Fig. S2A**). Moreover, only *MDK* expression showed a significant positive correlation with both our well-stablished pluripotency gene set and the *ALK* OE signature mentioned above (**Fig. S2B**). These results obtained in bulk tumor samples were confirmed in a PDAC single-cell transcriptomic dataset^[35]^, where *MDK* showed the strongest positive correlation with three out of the four of the stemness genes separately (*KLF4*=0.37, *OCT3/4*=0.9, *NANOG*=0.78, *SOX2*=0.81) (**Fig. 2A**).Further analyses of this single-cell dataset revealed that *MDK* was expressed by a wide range of cell types, including ductal, acinar and tumor cells as well as fibroblasts (**Fig. 2B**), whereas *PTN* was mainly expressed by stromal cells (**Fig. 2B**). Moreover, these analyses further corroborated the low expression of *ALK* mRNA in PDAC tumors, as it was undetectable at single-cell level in the different PDAC cell populations included in this dataset (**Fig. 2B**).

**Figure 2.**
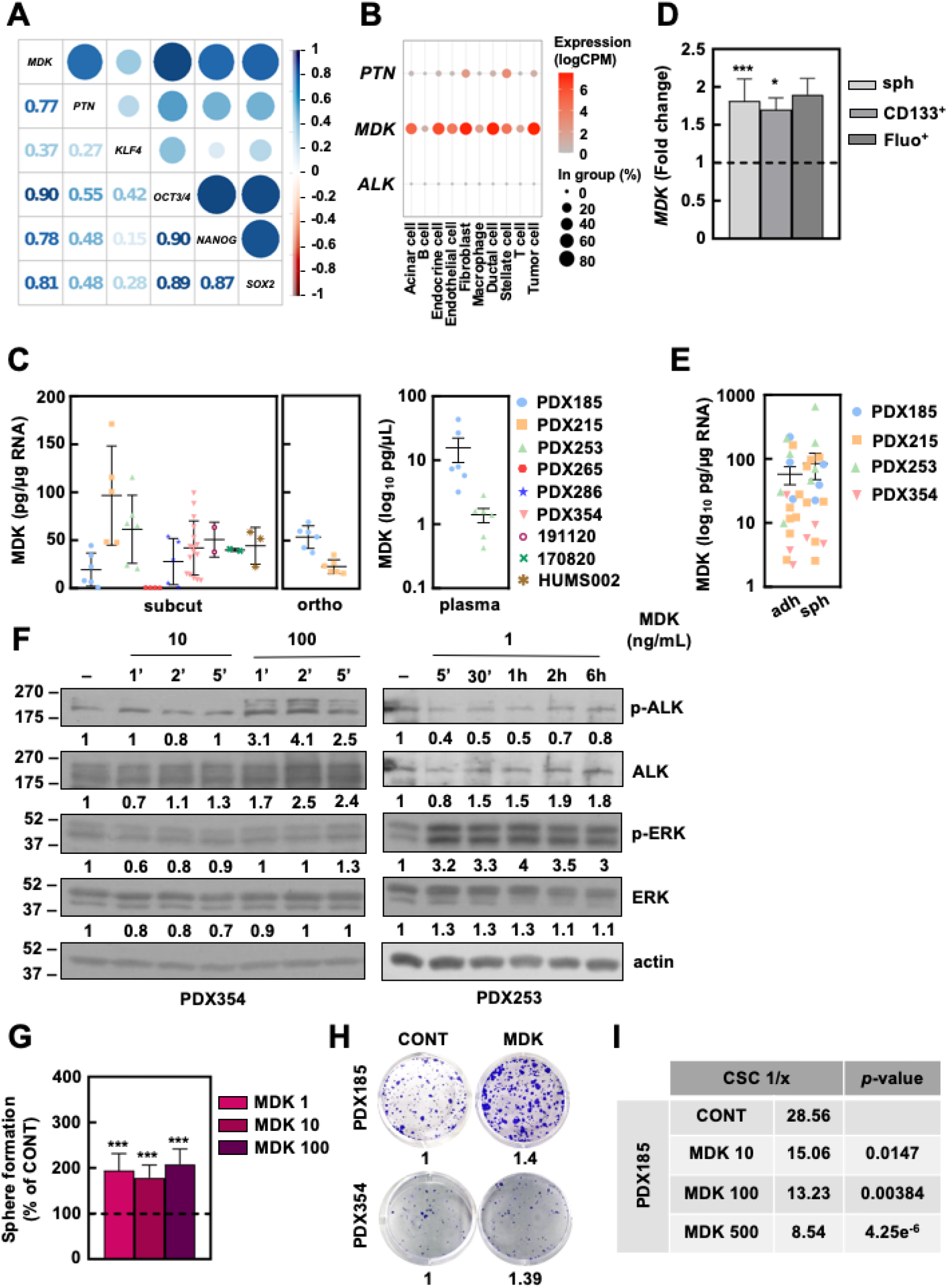
Ligand-dependent ALK activation supports self-renewal in PDAC. **A)** Correlation matrix of the indicated genes in single tumor cells from the Peng scRNAseq dataset. The numbers and dot sizes indicate the r value of each correlation. The blue color indicates a positive correlation, whereas the red color represents a negative correlation. **B)** Single-cell expression analysis of the indicated genes in the different cellular populations included in the Peng scRNAseq dataset. The dot size represents the percentage of cells expressing each gene per population, while the color scale denotes the expression level. **C)** MDK in supernatants from subcutaneous (SC, left panel) and orthotopic (O, middle panel) PDX implants *ex vivo* and in plasma (P, right panel) from orthotopic tumor-bearing mice by ELISA. **D)** RT-qPCR of *MDK* mRNA levels in the indicated CSC settings (sph: spheroids, sorted CD133^+^ and Fluo^+^; pooled data from PDX185 and 215). **E)** MDK detection in supernants from adherent (a) and spheroid (s) cultures by ELISA. **F)** Kinetics of ALK activation by western blot at the indicated times after treatment with 10 or 100 ng/mL of recombinant MDK. Numbers represent the quantification of the band intensity of each protein normalized by actin, shown as fold change from the control group. **G)** Sphere formation assay after pre-treatment with recombinant MDK for 72 hours at the indicated concentrations (ng/mL) in adherent conditions (pooled data from PDX185 and 354). The dashed lines represent the value of the differentiated cells (D) or control groups (G). Data are represented as mean ± SEM and analyzed using one-way ANOVA or Kruskal-Wallis tests of, at least, three independent experiments, unless otherwise specified. * p<0.05, ** p<0.01, *** p<0.005. **H)** Representative colony formation assay after 21 days of treatment with 10 ng/mL of recombinant MDK. The numbers represent the absorbance of crystal violet shown as the fold change from the control group. **I)** Estimation of the CSC frequency by *in vitro* extreme limiting dilution assay (ELDA) after treatment with the indicated concentrations of recombinant MDK for seven days. The numbers indicate one CSC every x number of cells.

Importantly, we confirmed that MDK was secreted by both subcutaneous and orthotopic PDX tumors *ex vivo* (**Fig. 2C, left and middle panel**, respectively) and *in vivo*, since we detected human MDK in the plasma of mice bearing orthotopic PDXs (**Fig. 2C, right panel**). Although CSC-enriched samples showed increased levels of *MDK* expression at the mRNA level (**Fig. 2D**), no significant differences between CSCs and differentiated cells were found in terms of MDK secretion (**Fig. 2E**). Treatment with recombinant human MDK induced ALK phosphorylation in the short term. Later on, phosphorylation was observed in its well-described downstream signaling partner ERK1/2^[36]^ (**Fig. 2F**), corroborating ligand-dependent ALK activation. This activation resulted in improved CSC functionality, as exogenous treatment with MDK enhanced self-renewal (**Fig. 2G**) and clonogenic capacity (**Fig. 2H**), and increased CSC frequency *in vitro* (**Fig. 2I**). Interestingly, similar results were obtained after treatment with PTN (**Fig. S2C-E**). These results confirm that ligand-dependent activation of the ALK pathway enhances PDAC aggressiveness by boosting CSCs properties.

#### *ALK inhibition abrogates CSC functionality* in vitro *and* in vivo

The use of small compounds, like Crizotinib or Ensartinib, to inhibit ALK signaling is a common approach to treat ALK^+^ malignancies, such as non-small lung cancer^[37]^. Considering ALK contribution to stemness in PDAC, we decided to test the effects of these compounds on our PDXs, including a model of metastatic PDAC established from circulating tumor cells (CTCA).

First, both Crizotinib and Ensartinib inhibited cell proliferation, with IC_50_ ranging from 0.7 to 3.8 and 0.4 to 1.8 µ M, respectively (**Fig. 3SA**). Importantly, both compounds inhibited ALK phosphorylation and downstream signaling at the selected concentrations (**Fig. 3A**). Since ALK inhibition with Crizotinib induced cell toxicity (**Fig. S3B**), we measured next cell death after treatment with both compounds. Treatment with both Crizotinib and Ensartinib induced apoptosis in the tumor bulk (**Fig. S3C, S3D**) and, most importantly, in the CD133^+^ population (**Fig. 3B, 3C**), thus decreasing the CD133 content (**Fig. 3D, 3E**).

**Figure 3.**
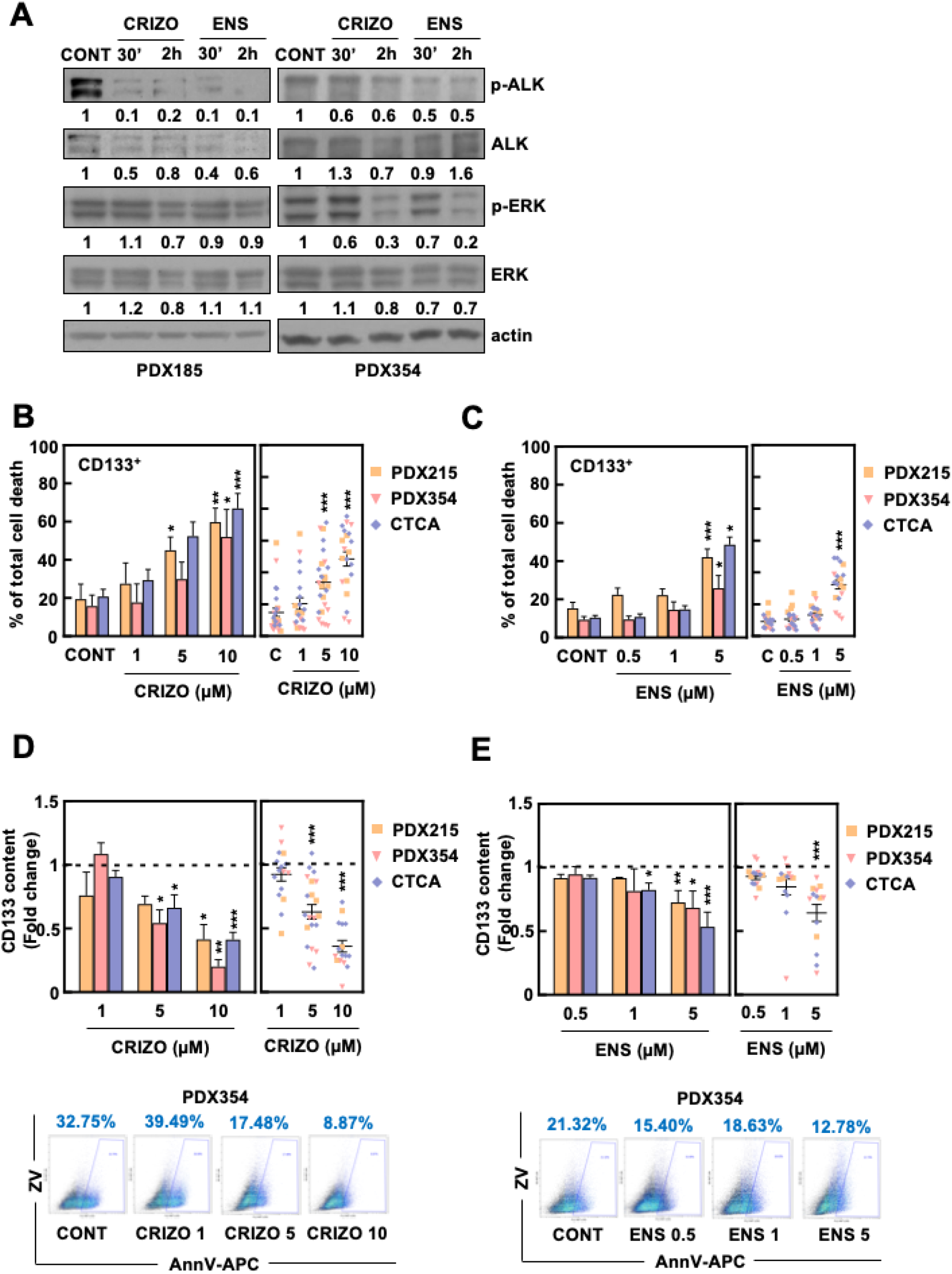
ALK pharmacologic inhibition targets PaCSCs. **A)** Kinetics of ALK inhibition after the indicated times of treatment with 10 µM Crizotinib and 5 µM Ensartinib. The numbers represent the quantification of the band intensity of each protein normalized by actin, shown as the fold change from control group. **B, C)** Percentage of total cell death measured as the sum of Annexin V^+^, Zombie Violet^+^ and double positive staining in CD133^+^ cells after 48 hours of treatment with Crizotinib (B) and Ensartinib (C). Left panels: mean value of each PDX separately. Right panels: pooled data, showing the individual values of each PDX. **D, E)** CD133 content in samples from B (D) and C (E). Top panels: mean of the CD133^+^ content of each PDX separately (left) and pooled data showing the individual values of each PDX (right); bottom panels: representative flow cytometry density plots of PDX354. Data are shown as the fold change from control group, which is represented with the dashed line. Data are represented as mean ± SEM and analyzed using one-way ANOVA or Kruskal-Wallis test of, at least, three independent experiments. * p<0.05, ** p<0.01, *** p<0.005.

Afterwards, since these findings suggested that ALK inhibition particularly targets PaCSCs, we assessed the efficacy of these compounds in impairing stemness-related functionality. Indeed, both Crizotinib and Ensartinib diminished self-renewal.(**Fig. 4A and S4A**) and clonogenic capacity (**Fig. 4B and S4C)**. Likewise, pretreatment with both Crizotinib and Ensartinib treatment decreased CSC frequency *in vitro* (**Fig. 4C and S4B**). These effects on CSC functionality could be validated *in vivo*, where pretreatment with the compounds decreased the number and size of tumors (**Fig. 4D and S4D**), the percentage of tumorigenicity (**Fig. 4E**) and the CSC frequency (**Fig. 4F**). Importantly, even though the percentage of epithelial tumor cells remained unchanged (**Fig. S4E**), the number of CD133^+^ cells decreased in the tumors obtained from Crizotinib-pretreated cells (**Fig. S4F**). These results demonstrate that ALK inhibition targets PaCSCs, by inducing cell death and effectively impairing their functionality.

**Figure 4.**
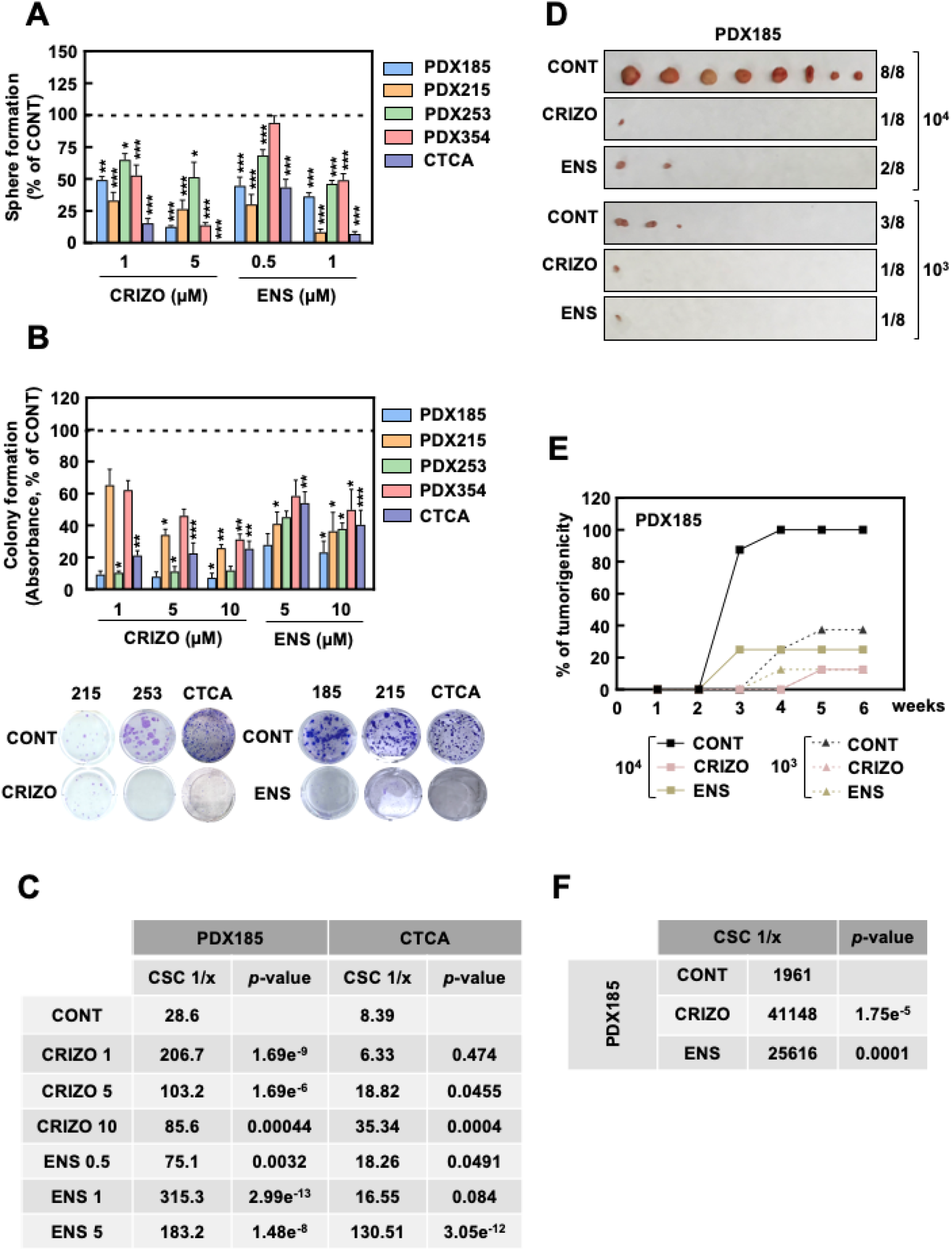
ALK pharmacologic targeting abrogates CSC features *in vitro* and *in vivo*. **A)** Sphere formation assay after seven days of treatment with Crizotinib and Ensartinib. **B)** Colony formation assay after 21 days of treatment with Crizotinib and Ensartinib. Top panel: absorbance of crystal violet. Bottom panel: images of a representative experiment of each PDX treated with either 1 μM Crizotinib or Ensartinib. **C)** CSC frequency after treatment with Crizotinib and Ensartinib for seven days, estimated by *in vitro* extreme limiting dilution assay (ELDA). **D-F)** *In vivo* ELDA of cells pre-treated *in vitro* with 10 µM Crizotinib and 5 µM Ensartinib for 48 hours and subcutaneously injected into the flanks of nude mice at decreasing cell densities. **D)** Pictures of tumors at end point (week six). **E)** Percentage of tumorigenicity over time. Note that the curves representing the Crizotinib conditions 10^3^ and 10^4^ cells overlap. **F)** Estimated CSC frequency. Data are represented as mean ± SEM and analyzed using one-way ANOVA or Kruskal-Wallis tests of, at least, three independent experiments. * p<0.05, ** p<0.01, *** p<0.005.

### *ALK inhibition prevents chemoresistance* in vitro *and* in vivo

One of the main contributors to chemotherapy failure in PDAC is its intrinsic chemoresistance^[38]^. Indeed, conventional chemotherapy usually targets just the tumor bulk, thus enriching the content of chemoresistant CSCs and causing tumor relapse. Considering the toxic effect of ALK inhibitors on PaCSCs, we decided to test if either Crizotinib or Ensartinib sensitized to Gemcitabine treatment *in vitro*. Interestingly, the combination of Gemcitabine with Crizotinib and Ensartinib independently at low doses decreased considerably the IC_50_ for this chemotherapeutic agent (**Fig. 5A and S5A**), resulting in enhanced cell death (**Fig. S5B**) Certainly, co-treatment with Ensartinib significantly increased cell death in the CD133 population (**Fig. 5B**) and decreased CD133 content (**Fig. 5C**) when compared to Gemcitabine alone. Importantly, both Crizotinib and Ensartinib prevented Gemcitabine-induced self-renewal (**Fig. 5D**) and clonogenic (**Fig. S5C**) capacity, indicating the effectiveness of this combined treatment.

**Figure 5.**
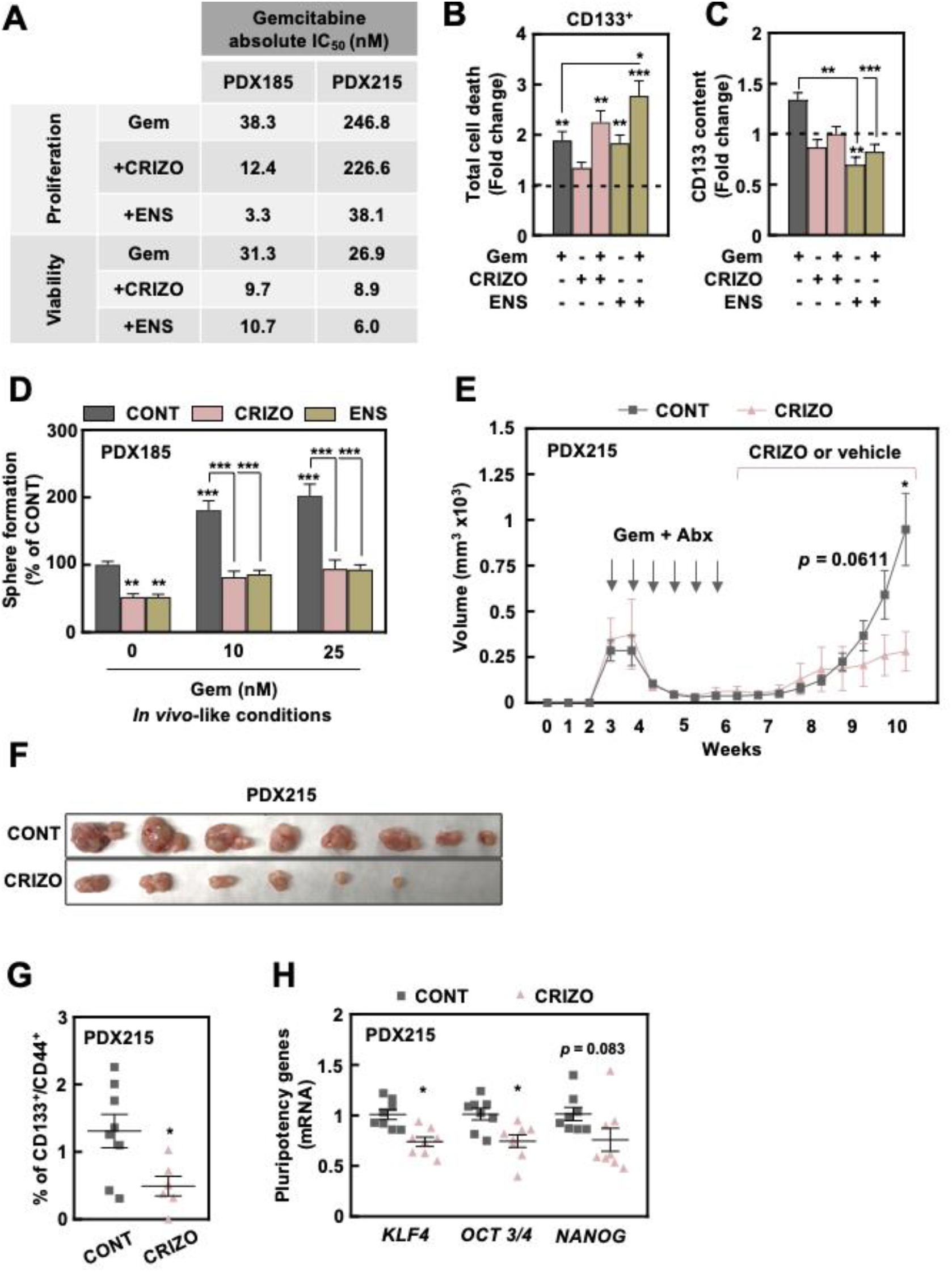
ALK inhibition synergizes with Gemcitabine treatment *in vitro* and *in vivo*. The combined effect of Gemcitabine and ALK inhibitors was studied using low doses of the compounds (Gem 50 nM unless otherwise specified, Crizotinib 1 µM, Ensartinib 1 µM). **A)** IC_50_ of Gemcitabine at 72 hours of treatment alone and in combination with Crizotinib or Ensartinib. **B)** Total cell death measured as the sum of Annexin V^+^, Zombie Violet^+^ and double positive staining in CD133^+^ cells after 48 hours of treatment as indicated (pooled data from PDX185, 253 and 354). **C)** CD133 content of the samples shown in B. Data are shown as the fold change from control group, which is represented with a dashed line (B and C). **D)** Sphere formation assay mimicking the *in vivo* conditions: 48 hours of Gemcitabine pre-treatment in adherent conditions prior to seven days of treatment with Crizotinib and Ensartinib in anchorage-independent conditions as indicated. **E-H)** *In vivo* treatment of mice subcutaneously implanted with PDX215 tumour pieces. When tumors reached around 300 mm^3^, mice were treated with 30 mg/kg of Abraxane (i.v., twice a week) in combination with 70 mg/kg of Gemcitabine (i.p., once a week) for 21 days. After seven days of rest, mice were randomized and treated with either vehicle or 25 mg/kg of Crizotinib (oral gavage, twice a day) until end point. **E)** Tumor volume over time. **F)** Pictures of tumors at end point (week 10). **G)** Percentage of CD133^+^/CD44^+^ cells of tumors shown in F. **H)** Pluripotency gene expression of tumors shown in F. Data are represented as mean ± SEM and analyzed using one-way ANOVA or Kruskal-Wallis tests of, at least, three independent experiments. * p<0.05, ** p<0.01, *** p<0.005

Considering these results, we decided next to translate this approach into the *in vivo* setting, by treating mice bearing subcutaneous PDAC implants with Crizotinib after a chemotherapy cycle with Gemcitabine and Abraxane, the most commonly used chemotherapy combination to treat PDAC nowadays. We first confirmed that treatment with Crizotinib was not toxic to the animals by showing that their body weight remained essentially unchanged (**Fig. S5D**). Strikingly, treatment with Crizotinib significantly delayed tumor growth after chemotherapy (**Fig. 5E, 5F**). Crizotinib-treated tumors were significantly smaller (**Fig. S5E**) and lighter (**Fig. S5F**) than tumors in the control group, and in some cases disappeared completely after treatment (**Fig. 5F**). Importantly, while no difference was found in the expression of the epithelial marker EpCAM (**Fig. S5G**), Crizotinib-treated tumors showed decreased content of CD133^+^/CD44^+^ cells (**Fig. 5G**) and reduced expression of stemness genes (**Fig. 5H**). These results confirmed our hypothesis that blocking ALK using small molecule inhibitors delays tumor growth after chemotherapy by targeting PaCSCs.

Taken together, our findings demonstrate an important role of ALK receptor in PaCSCs contributing to PDAC aggressiveness. Importantly, the use of chemotherapeutic agents in combination with ALK inhibitors shows the potential for mitigating the otherwise inevitable tumor relapse after chemotherapy, thus improving treatment outcome in PDAC patients.

## DISCUSSION

Despite current efforts to improve treatment outcomes, life expectancy of pancreatic cancer remains terribly brief. This is, at least in part, due to the presence of aggressive pancreatic cancer stem cells (PaCSCs) that survive after conventional chemotherapy, eventually regrow the tumor and migrate to colonize secondary organs. For this reason, targeting CSCs in combination with conventional therapies may be the only way to ensure long-term survival of pancreatic ductal adenocarcinoma (PDAC) patients.

Previous studies from our group and others, have shown that PaCSCs bear unique features, essential to maintain their properties and functionality and, in principle, are amenable for therapeutic intervention. Indeed, we have provided proof-of-concept for the efficacy of metabolic inhibition for PaCSCs targeting in animal models^[33,39–41]^, but further clinical translation has remained challenging due to lack of clinically effective compounds. In contrast, receptor and non-receptor tyrosine kinases (RTKs and nRTKs, respectively) represent a much more approachable strategy since they can be targeted by a plethora of specific and clinical-grade compounds. In addition, RTKs and nRTKs control essential cellular mechanisms dysregulated in cancer, such as metabolism, proliferation, survival and, most importantly, stemness^[11,12,14,15]^. Several clinical trials in PDAC are currently exploring the potential benefit of diverse TK inhibitors in PDAC^[42]^.

In order to find novel pharmacologically amenable targets hyperactivated in PaCSCs, we decided to focus on RTKs, since their expression in the cellular membrane allows for targeting by both small molecules and blocking antibodies. Screening of RTKs revealed that the receptor anaplastic lymphoma kinase (ALK) was consistently overexpressed and hyperactivated in PaCSCs, using different CSC enrichment methods and PDAC patient-derived xenograft (PDX) models (**Fig. 1A, 1B and S1A, S1C**) to account for intrinsic intra- and intertumoral heterogeneity. Although the ALK receptor plays an important physiological role in neural development^[26,43]^, it was first discovered in lymphoma as the fusion protein NPL-ALK following chromosomal rearrangement^[44]^. Subsequently, ALK was shown to be aberrantly expressed and/or activated in several cancer types^[26–29]^. ALK receptor activation triggers different intracellular signaling pathways involved in proliferation, survival and metabolism, including JAK/STAT and Ras/ERK^[36,45]^. Importantly, some studies suggested that ALK acts as a regulator of stemness in several cancers^[46–49]^. However, our results were certainly unexpected, since this receptor has been overlooked in PDAC. The possible cause points to the lack of mRNA overexpression in tumor bulk cells when compared with normal pancreas or the low level of chromosomal rearrangement of the *ALK* gene in PDAC^[30]^ (**Fig. S1D, S1E**), which is the main pathogenic mechanism associated to ALK in NSCLC and brain tumors^[26–29]^. The low *ALK* expression at the mRNA level detected in bulk transcriptomic analysis of The Cancer Genome Atlas (TCGA) dataset (**Fig. S1D**) was further evidenced in the single-cell expression analysis (scRNAseq, **Fig. 2B**), where *ALK* was undetectable in the populations of the pancreatic niche, including tumor cells. In fact, detection of *ALK* mRNA levels proved challenging even in our primary cultures (**Fig. S1C and data not shown**). However, we detected considerable ALK expression and phosphorylation in different PDAC PDX models and established cell lines by western blot (**Fig. 1B, 1C and S1C**). Our data further reinforces the importance of considering protein post-translational regulation and modifications over purely transcriptomic studies for target discovery screenings.

Nevertheless, further bioinformatic analyses of transcriptomic datasets supported our initial results *in vitro*. Indeed, we were able to link the mRNA expression of both *ALK* and an *ALK* overexpression (OE) signature previously described^[23]^ with pathways related with CSCs in PDAC patients, such as epithelial-to-mesenchymal transition (EMT), drug metabolism and stemness (**Fig. 1D, 1E**). In addition, these analyses allowed us to propose the cytokines midkine (MDK) and, to a lesser extent, pleiotrophin (PTN), as the main putative ligands triggering ALK activation in PaCSCs. Indeed, the identification of the actual ALK ligand is a matter of great controversy: while some studies point to MDK and PTN^[27,50]^, others suggest the cytokines FAM150A and B (family with sequence similarity 150 members A and B)^[34,51]^ as main activators of the ALK pathway. While the results obtained for *FAM150A* and *FAM150B* were inconsistent, both *MDK* and *PTN* were overexpressed in PDAC patient samples, although only *MDK* correlated with the expression of pluripotency genes in both bulk and scRNAseq data and with the *ALK* OE signature mentioned above (**Fig. 2A and S2A-C**).

MDK and PTN are heparin-binding growth factors with multiple regulatory functions in biological processes such as proliferation, differentiation and development through binding to different receptors, including ALK^[52,53]^. Interestingly, functional assays with recombinant MDK and PTN demonstrated that ligand-dependent ALK activation increased self-renewal, clonogenicity and CSC frequency in our PDX models (**Fig. 2F-I and S2D-F**), indicating that the axis MDK/PTN-ALK enhances stemness in PDAC. Indeed, ALK activation *via* MDK or PTN has been shown to regulate self-renewal and tumorigenicity in glioblastoma^[46,47]^, while PTN knockdown favored chemosensitivity and inhibited clonogenic capacity in osteosarcoma^[54]^.

Our results point to different modes of ALK ligand-dependent activation in PaCSCs. On the one hand, *PTN* was barely expressed in tumor cells according to our analysis of the scRNAseq PDAC dataset (**Fig. 2B**) and was undetectable in PDXs (data not shown). In contrast, *PTN* was expressed by stromal cells, such as fibroblasts and stellate cells (**Fig. 2B**), revealing a potential paracrine regulatory loop in which cells from the tumor microenvironment may sustain PaCSC through ALK activation *via* PTN. On the other hand, analysis of the scRNAseq dataset indicated that *MDK* was expressed by a wide range of cells present in the pancreatic niche, including tumor cells (**Fig. 2A, 2B**). We confirmed high levels of human MDK secretion in supernatants of both subcutaneous and orthotopic tumor pieces, plasmas from orthotopic tumor-bearing mice (**Fig. 2C**) and primary tumor cells in culture regardless of their pluripotency status (**Fig. 2E**). Strikingly, a recent study revealed that melanoma cells secrete MDK to promote an immune-suppressive microenvironment involving tumor associated macrophages and cytotoxic T cells^[55]^, suggesting that MDK secretion by PaCSCs could also play a role in immunoediting in PDAC.

Importantly, we have demonstrated the crucial role of ALK for PDAC stemness not only by exogenous activation of the receptor, but also through pharmacological inhibition strategies using clinically-approved compounds. Crizotinib is a trivalent ALK, c-Met and ROS1 inhibitor approved by the Food and Drug Administration (FDA) to treat cancers expressing oncogenic ALK fusion proteins^[56,57]^ and, later on, those depending on c-Met and/or ROS1 signaling^[58]^. On the other hand, Ensartinib is a potent and specific next-generation ALK inhibitor currently evaluated in a phase III trial^[59]^, included in our study to discard off-target effects induced by Crizotinib. Treatment with both Crizotinib and Ensartinib inhibited proliferation (**Fig. S3A**), and induced cell death *in vitro* (**Fig. 3B, 3C and S3B, S3C**), suggesting that ALK signaling contributes to cell survival. ALK inhibition was effective in our PDX models derived from both local (PDXs) and metastatic PDAC (CTCA), suggesting this therapeutic approach may be effective in advanced and metastatic patients. Notably, ALK inhibition drastically decreased the CSC content (**Fig. 3D**) and stemness features *in vitro* and *in vivo* (**Fig. 4 and S4**), demonstrating that ALK is functionally necessary for PaCSC maintenance.

Chemotherapy failure remains a major issue in PDAC management due to its intrinsic chemoresistance. In addition, conventional treatments target the tumor bulk and enriches the CSC content, responsible for tumor relapse. Here, we demonstrate that both Crizotinib and Ensartinib decreased the IC_50_ of Gemcitabine more than half (**Fig. 5A and S5A**) with stronger effects by Ensartinib treatment. Importantly, both ALK inhibitors in combination with Gemcitabine decreased PaCSC content (**Fig. 5B, 5C**) and abrogated Gemcitabine-induced self-renewal (**Fig. 5D**) and clonogenicity (**Fig. S5C**). Importantly, Crizotinib treatment significantly delayed tumor relapse *in vivo* (**Fig. 5E**) and some tumors even disappeared after treatment (**Fig. 5F**). Moreover, tumors treated with Crizotinib showed reduced stemness markers (**Fig. 5G, 5H**), indicating a successful PaCSC targeting *in vivo*. Since development of resistance to Crizotinib has been reported already^[60]^, further *in vivo* experiments would be needed in order to test Ensartinib, a compound still in the process of approval by the FDA. Besides its improved specificity, this inhibitor was more potent than Crizotinib either alone or in combination with Gemcitabine *in vitro*. Considering our results, we would expect a reduction of the required dosage *in vivo* to obtain a positive response, further translated into minimal side effects when applied as combinatory treatment.

In summary, our results demonstrate that PaCSCs sustain their stemness program through MDK –and PTN–-dependent activation of the ALK signaling pathway. Importantly, this pathway can be pharmacologically targeted with small molecule inhibitors that, combined with conventional chemotherapy, show promising effects for an effective long-term treatment of PDAC.

## AUTHOR CONTRIBUTIONS

**B. Parejo-Alonso**: conceptualization, investigation, formal analysis, visualization, writing-original draft, writing-review and editing. **A. Royo-García**: investigation, writing-review and editing. **P. Espiau-Romera**: investigation, writing-review and editing. **S. Courtois**: investigation, writing-review and editing. **A. Curiel-García**: formal analysis, visualization, writing-review and editing. **S. Zagorac**: investigation, writing-review and editing. **I. Villaoslada**: investigation, writing-review and editing. **K. P. Olive**: resources, writing-review and editing. **C. Heeschen**: resources, writing-review and editing. **P. Sancho**: conceptualization, project administration, supervision, funding acquisition, investigation, writing-original draft, writing-review and editing.

## ACKNOWLEDGEMENTS

Authors would like to acknowledge the use of the CIBA (Centro de Investigación Biomédica de Aragón) Flow Cytometry, Pathology and Microscopy Facilities (Servicios Científico-Técnicos, IACS-Universidad de Zaragoza). We also thank Laura Sancho Andrés for proofreadin the manuscript.The research was supported by the Instituto de Salud Carlos III through the Miguel Servet Program (CP16/00121 to P.S.), a PFIS predoctoral contract (FI21/00031 to P. E-R) and Fondo de Investigaciones Sanitarias (PI17/00082 and PI20/00921, to P.S.) (all co-financed by European funds (FSE: “El FSE invierte en tu futuro” and FEDER: “Una manera de hacer Europa”, respectively), the Worldwide Cancer Research (WCR) Charity together with Asociación Española contra el Cáncer (AECC) (19-0250, to P.S.). BP-A was supported by crowdfunding through Precipita-Fecyt and a predoctoral contract from Apadrina la Ciencia-Ford Motor Company. We are grateful to patients with pancreatic cancer who donated samples to the Barts Pancreatic Tissue Bank (http://www.bartspancreastissuebank.org.uk) funded by the Pancreatic Cancer Research Fund.

