## supplementary figures 1-5 for "ALK signaling drives tumorigenicity and chemoresistance of pancreatic ductal adenocarcinoma cells"

**Figure S1: ALK is preferentially activated and overexpressed in PaCSCs. A)** Mean of the quantification of the dots corresponding to p-ALK from Fig. 1A. DOP: density of pixels, Diff: differentiated, CSCs: cancer stem cells. **B)** Western blot of cell lysates comparing PDXs and cell lines. Numbers represent the quantification of the band intensity for each protein normalized by actin. Bx: BxPC3; Mia: MiaPaCa2; Su: Su8686. **C)** RT-qPCR for *ALK* mRNA levels in different CSC settings (pooled data from PDX185 and 215). Sph: spheroids. CD133 n=2, Fluo n=1. The dashed line represents the value of the differentiated cells. Data are represented as mean ± SEM and analyzed using one-way ANOVA or Kruskal-Wallis tests of, at least, three independent experiments, unless otherwise specified. * $p<0.05$, ** $p<0.01$, *** $p<0.005$. **D)** Transcriptomic bioinformatic analyses of *ALK* comparing normal (N) and PDAC (T) human tissues from TCGA and GTEx datasets. **E)** Mutational status of *ALK* from different datasets: 1. Pancreas UTSW, 2. Pancreas TCGA PanCan 2018, 3. Pancreas TCGA, 4. Pancreas ICGC and 5. Pancreas QCMG 2016. Mut: mutation, Amp: amplification, DDel: deep deletion.

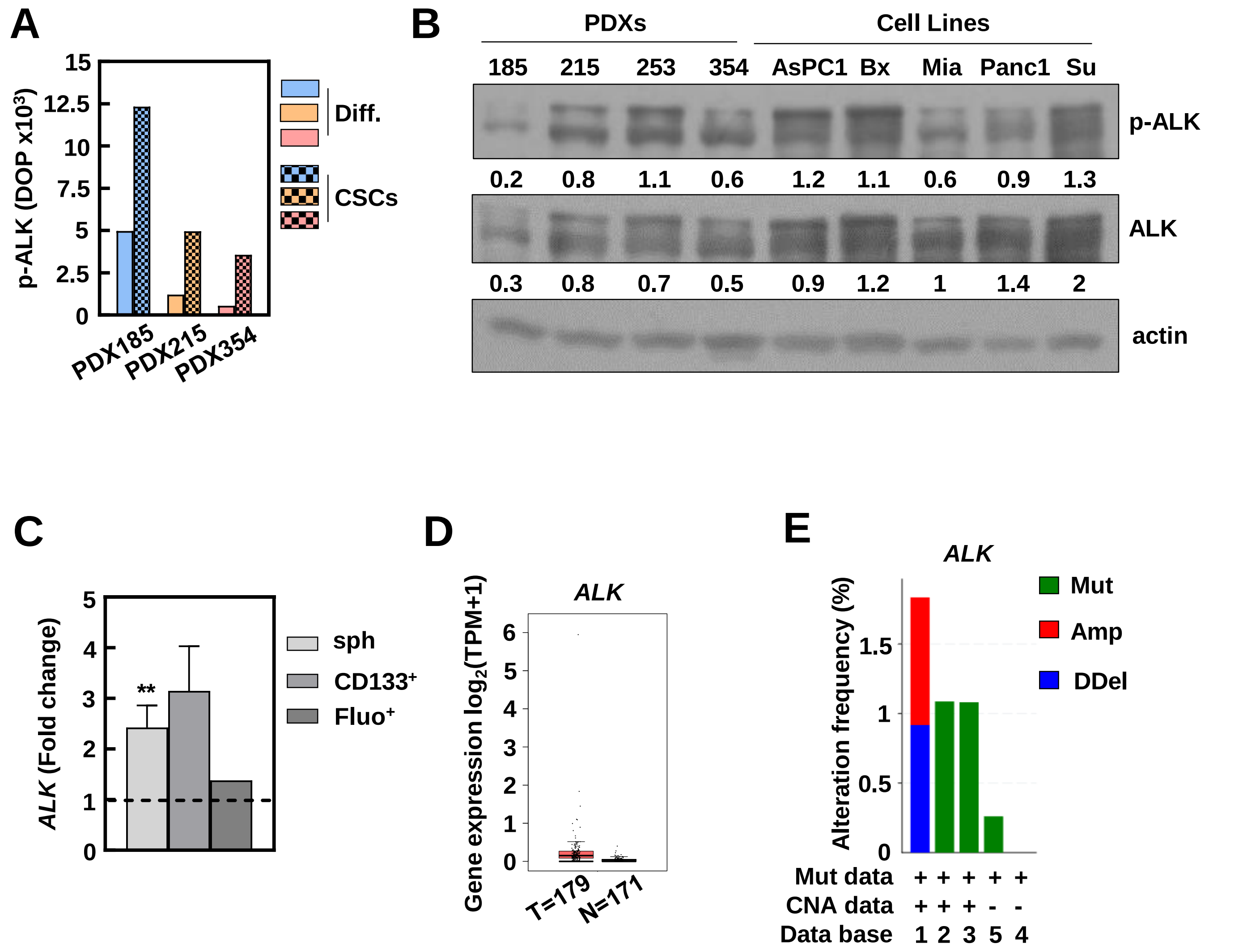

**Figure S2: Ligand-dependent ALK activation supports self-renewal in PDAC. A)** Expression of ALK ligands comparing normal (N) and PDAC (T) human tissues from TCGA and GTEx datasets. **B)** Correlation expression of ALK ligands with a pluripotency signature composed by *KLF4*, *OCT3/4*, *NANOG* and *SOX2* (top row) or an ALK overexpression (OE) signature (bottom row) in human tissues from TCGA and GTEx datasets. TPM: transcripts per million. **C)** Kinetics of ALK activation after the indicated times of treatment with 1 ng/mL of recombinant PTN measured by Western Blot. Numbers represent the quantification of the band intensity for each protein normalized by actin, shown as the fold change to control condition. **E)** Sphere formation assay after pre-treatment with recombinant PTN for 72 hours at the indicated concentrations (ng/mL) in adherent conditions (pooled data from PDX185 and 354). The dashed line represents the value of the control condition. Data are represented as mean ± SEM and analyzed using one-way ANOVA or Kruskal-Wallis tests of, at least, three independent experiments. * p<0.05, ** p<0.01, *** p<0.005. **F)** Representative colony formation assay after 21 days of treatment with 10 ng/mL recombinant PTN. The numbers represent the crystal violet staining absorbance, shown as the fold change to control condition.

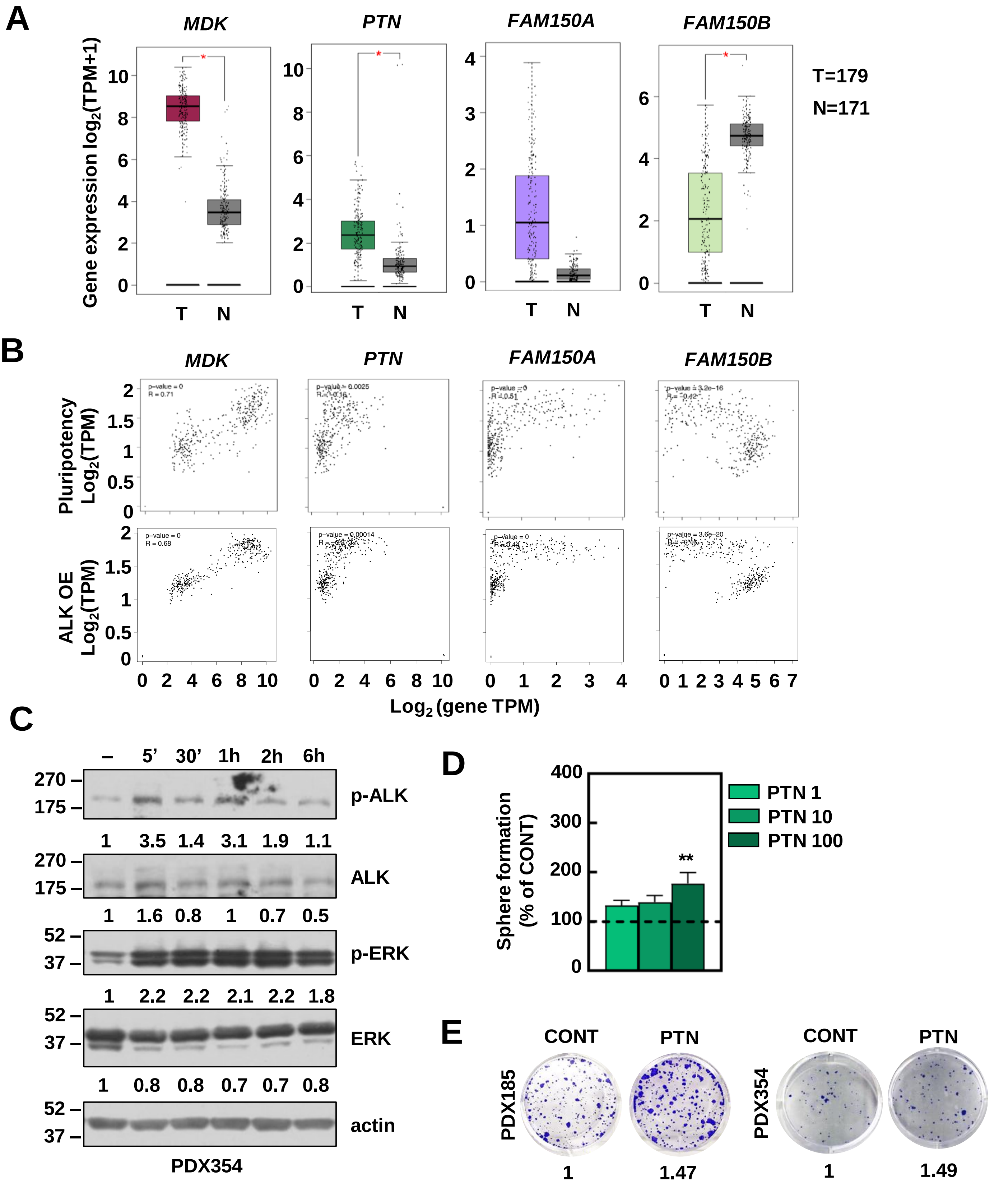

**Figure S3: ALK pharmacologic inhibition targets PaCSCs. A)** $IC_{50}$ of Crizotinib and Ensartinib at 72 hours of treatment. Left panel: $IC_{50}$ values for each cell type and inhibitor; right panel: representative proliferation rate graph of PDX185 and CTCA. **B)** Toxicity of Crizotinib for each cell type measured as relative fluorescence units normalized by crystal violet. Data are shown as the fold change to control condition which is represented as the dashed line (A and B). **C, D)** Percentage of total cell death measured as the sum of Annexin $V^+$, Zombie $Violet^+$ and double positive staining in the whole population after 48 hours of treatment with Crizotinib (C) or Ensartinib (D). Top panels: mean value for each PDX separately (left) and pooled data showing the individual values for each PDX (right); bottom row: representative flow cytometry density plots of PDX354. Data are represented as mean ± SEM and analyzed using one-way ANOVA or Kruskal-Wallis tests of, at least, three independent experiments. * p<0.05, ** p<0.01, *** p<0.005.

**A**

| | Absolute $IC_{50}$ (µM) | |
|---|---|---|
| | CRIZO | ENS |
| PDX185 | 3.745 | 1.760 |
| PDX215 | 0.809 | 0.3901 |
| PDX253 | 2.173 | 1.410 |
| PDX354 | 1.097 | 0.7847 |
| CTCA | 0.681 | 1.052 |

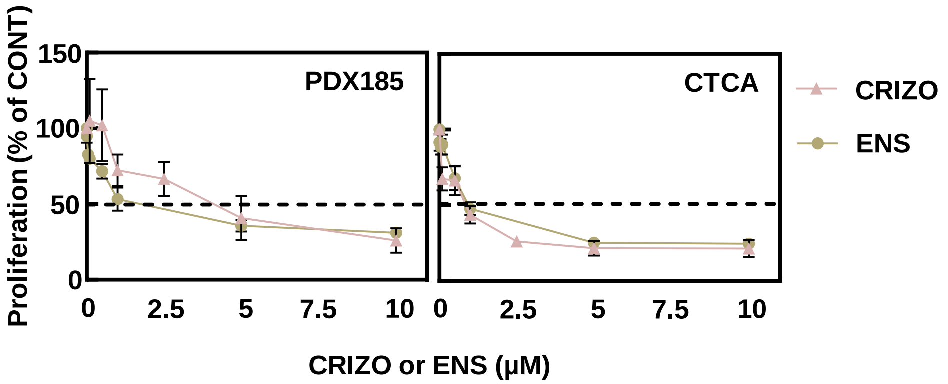


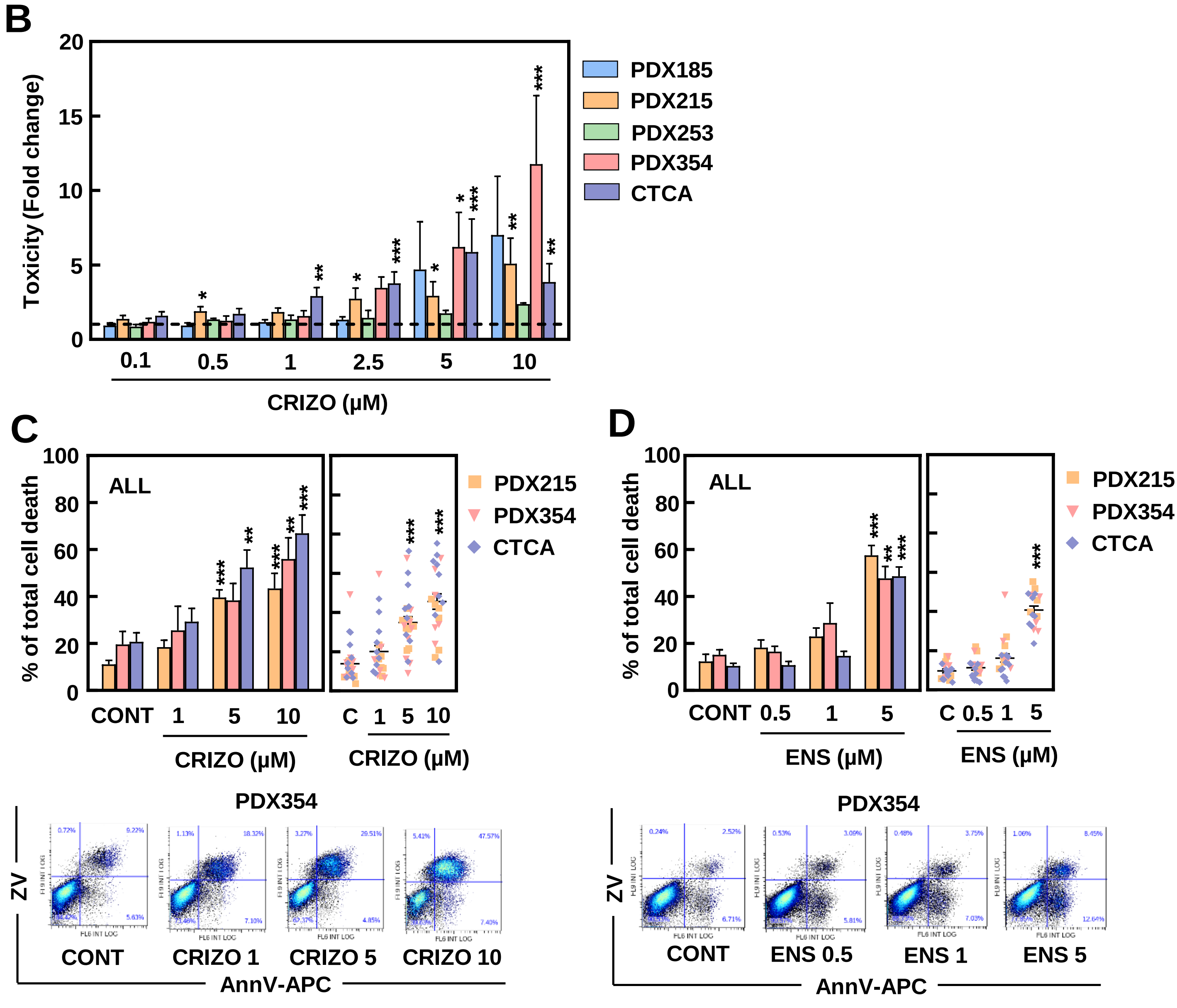

**Figure S4. ALK pharmacologic targeting abrogates CSC features *in vitro* and *in vivo*. A)** Sphere formation assay from Fig. 4A. Left panel: individual values for each PDX; right panel: representative images of PDX185. **B)** Sphere formation assay after pre-treatment with Crizotinib for 48 hours in adherent conditions. Left panel: percentage of spheres for each PDX; right panel: pooled data showing the individual values for each PDX. ND: not determined. **C)** Colony formation assay after 21 days of treatment with Crizotinib or Ensartinib. Pooled data showing the individual values for each PDX. Data are shown as the percentage of control condition which is represented as the dashed line. **D-F)** *In vivo* ELDA of cells pre-treated *in vitro* with 5 µM Crizotinib for 48 hours and subcutaneously injected in the flanks of nude mice at decreasing cell densities. **D)** Tumors at end point (week 6); **E)** percentage of $EpCAM^{+}$ cells in the $10^4$ tumors showed in D; **F)** $CD133^{+}$ content in the $10^4$ tumors showed in D. Data are represented as mean ± SEM and analyzed using one-way ANOVA or Kruskal-Wallis tests of, at least, three independent experiments. * p<0.05, ** p<0.01, *** p<0.005.

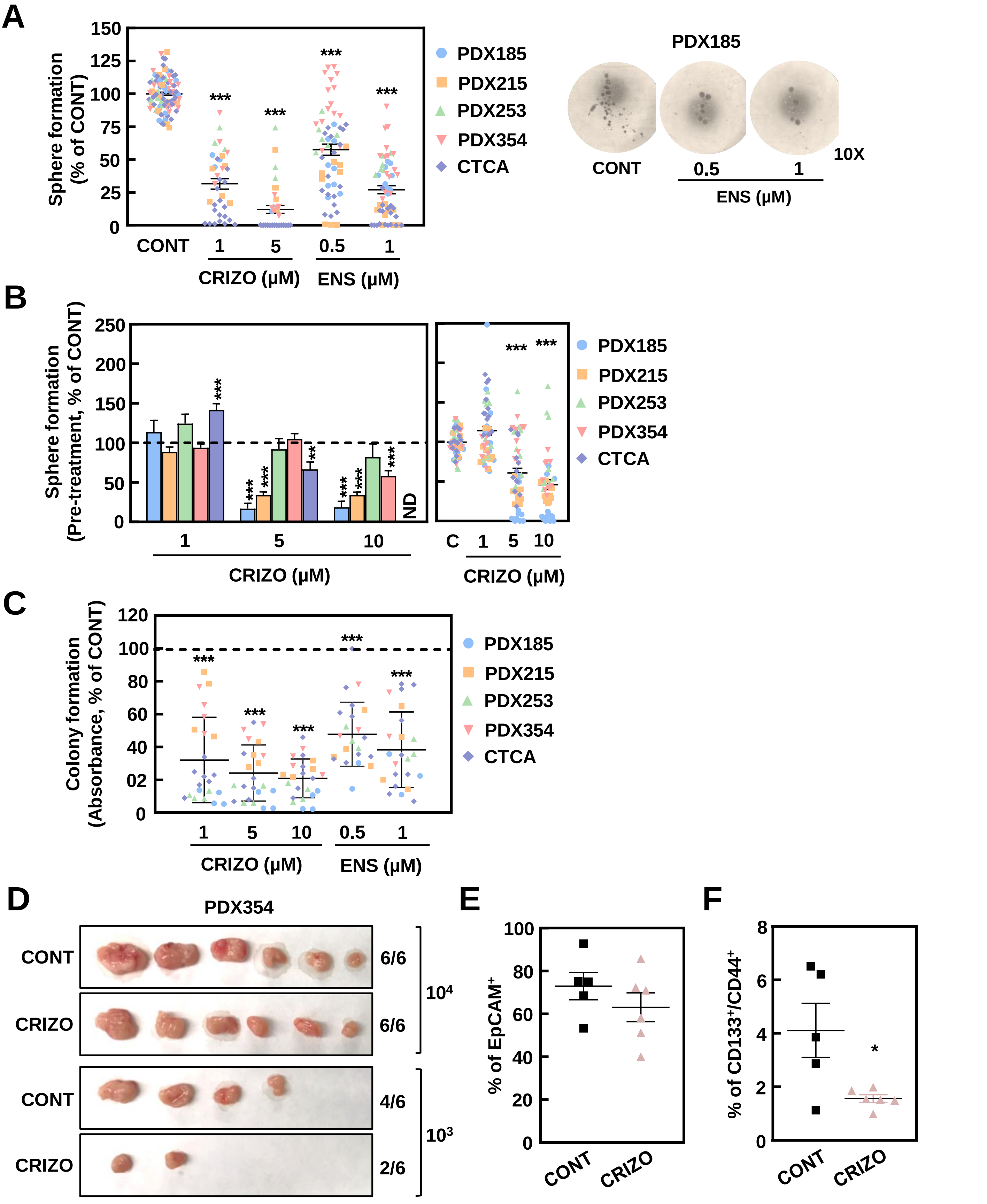

**Figure S5: ALK inhibition synergizes with Gemcitabine treatment *in vitro* and *in vivo*.** The combined effect of Gemcitabine and ALK inhibitors was studied using low doses of the compounds (Gem 50 nM unless otherwise specified, Crizotinib 1 µM, Ensartinib 1 µM). **A)** Representative proliferation rate graph of PDX185 and 215 from Fig. 5A. **B)** Total cell death measured as the sum of Annexin $V^+$, Zombie $Violet^+$ and double positive staining in the whole population after 48 hours of treatment with Gemcitabine alone or in combination with Crizotinib or Ensartinib (pooled data from PDX185, 253 and 354). **C)** Colony formation assay after 21 days of treatment with Gemcitabine alone or in combination with Crizotinib or Ensartinib. **D-G)** *In vivo* treatment from Fig. 5E-H. **D)** Mice weight over time; **E)** tumor volume at end point (week 10); **F)** tumor weight at end point (10 weeks); **G)** percentage of $EpCAM^+$ cells of tumors shown in Fig. 5F. Data are represented as mean ± SEM and analyzed using one-way ANOVA or Kruskal-Wallis tests of, at least, three independent experiments. * p<0.05, ** p<0.01, *** p<0.005.

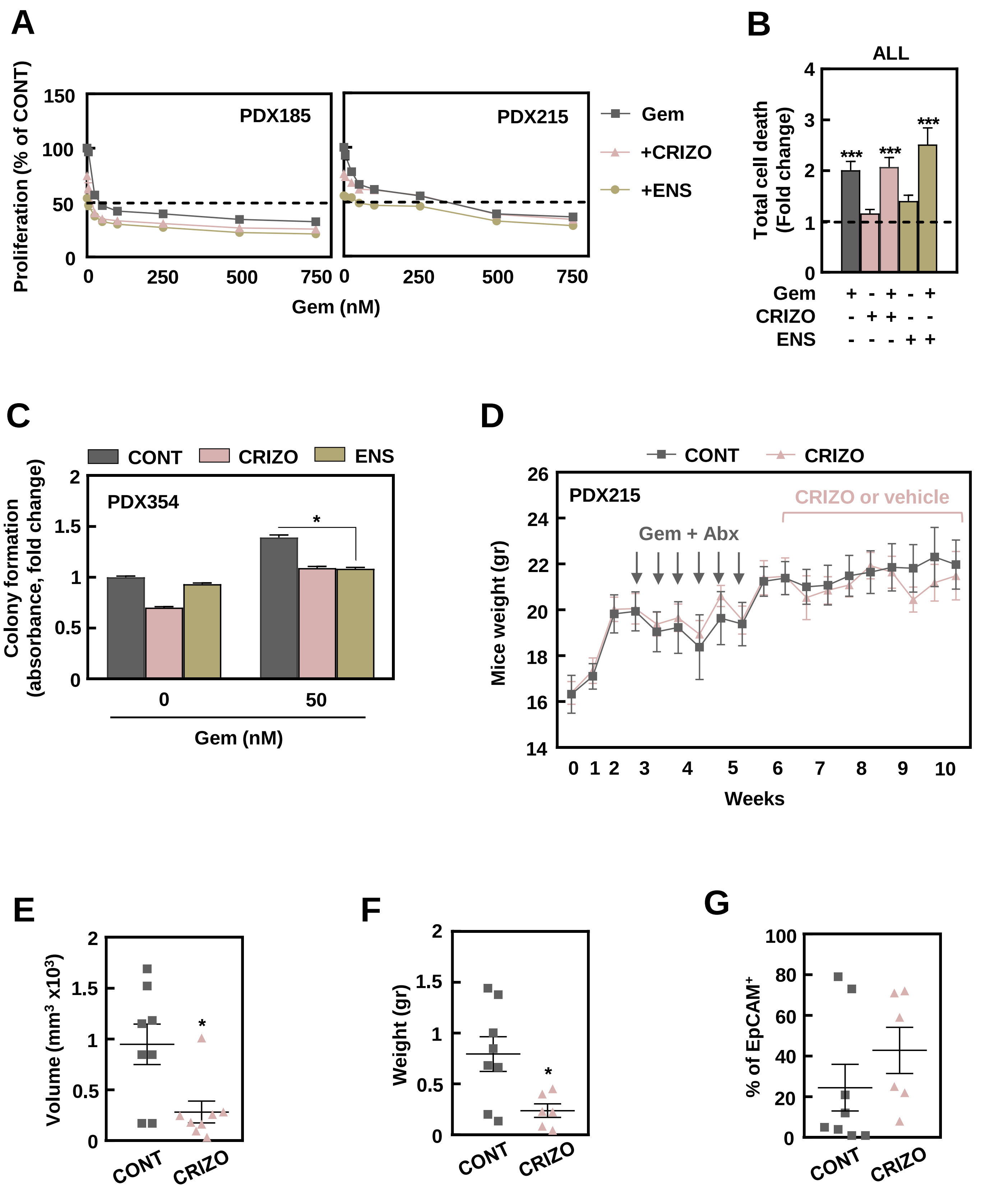
